## Supplementary figures and images for "MftG is crucial for ethanol metabolism of mycobacteria by linking mycofactocin oxidation to respiration"

### Supplementary Figure S1

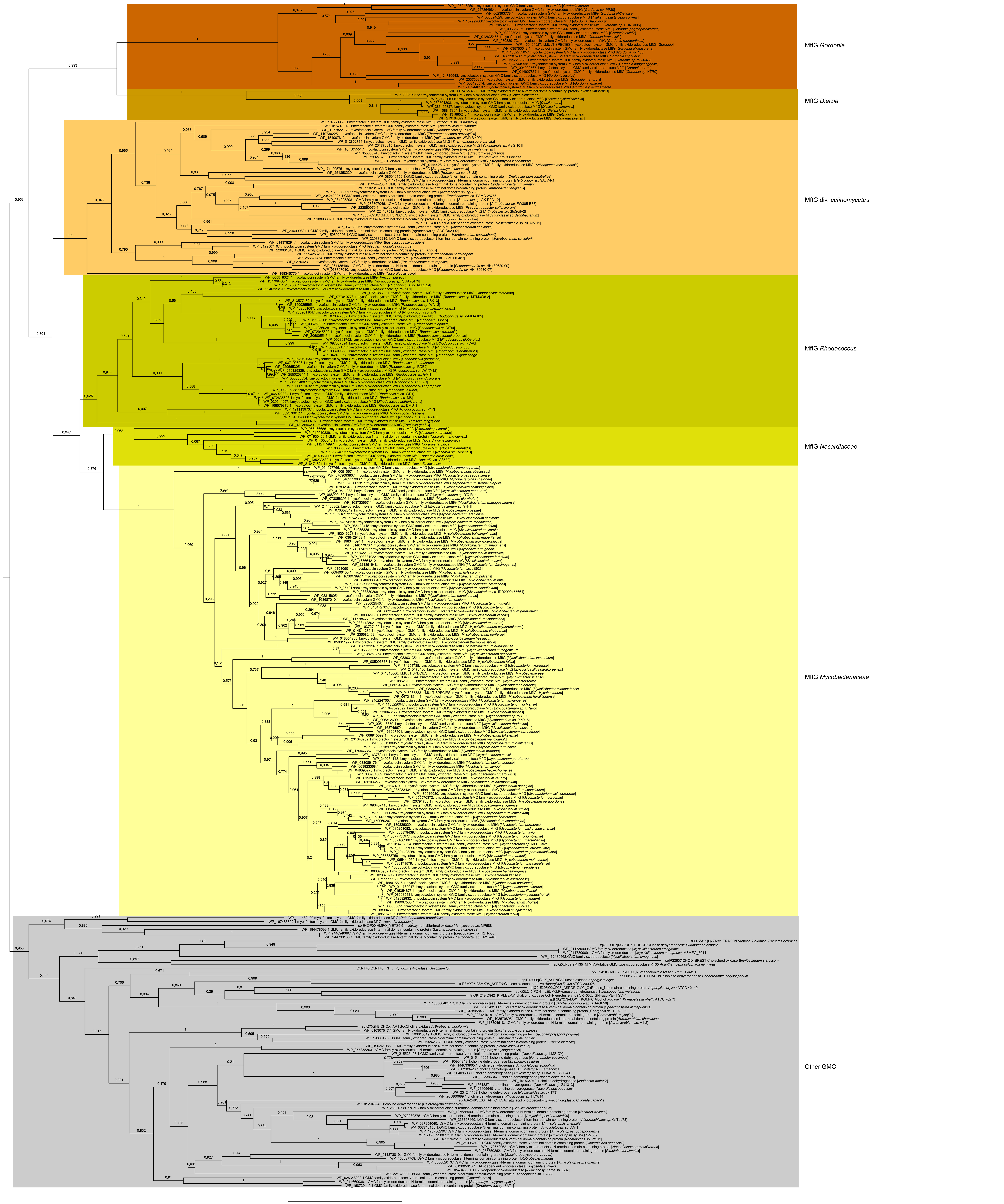
