## Supplementary Information for "MftG is crucial for ethanol metabolism of mycobacteria by linking mycofactocin oxidation to respiration"

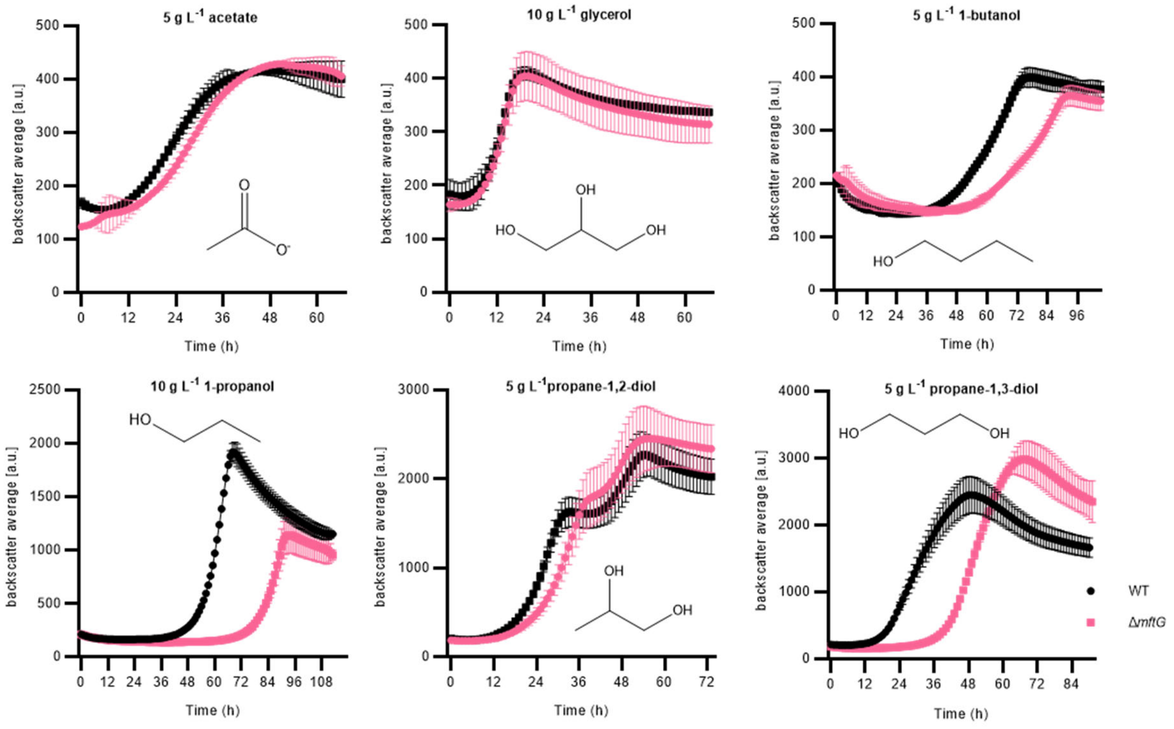


**Supplementary Figure S2 – Effect of *mftG* gene deletion on mycobacterial growth using different carbon sources**. Growth curve of *M. smegmatis* WT and Δ*mftG* growing on HdB-Tyl supplemented with 5 g $L^{-1}$ acetate, 10 g $L^{-1}$ glycerol, 10 g $L^{-1}$ 1-propanol, 5 g $L^{-1}$ 1-butanol, 5 g $L^{-1}$ propane-1,2-diol and 5 g $L^{-1}$ propane-1,3-diol. Measurements were performed in at least biological duplicates (n≥2). Error bars represent standard deviations.

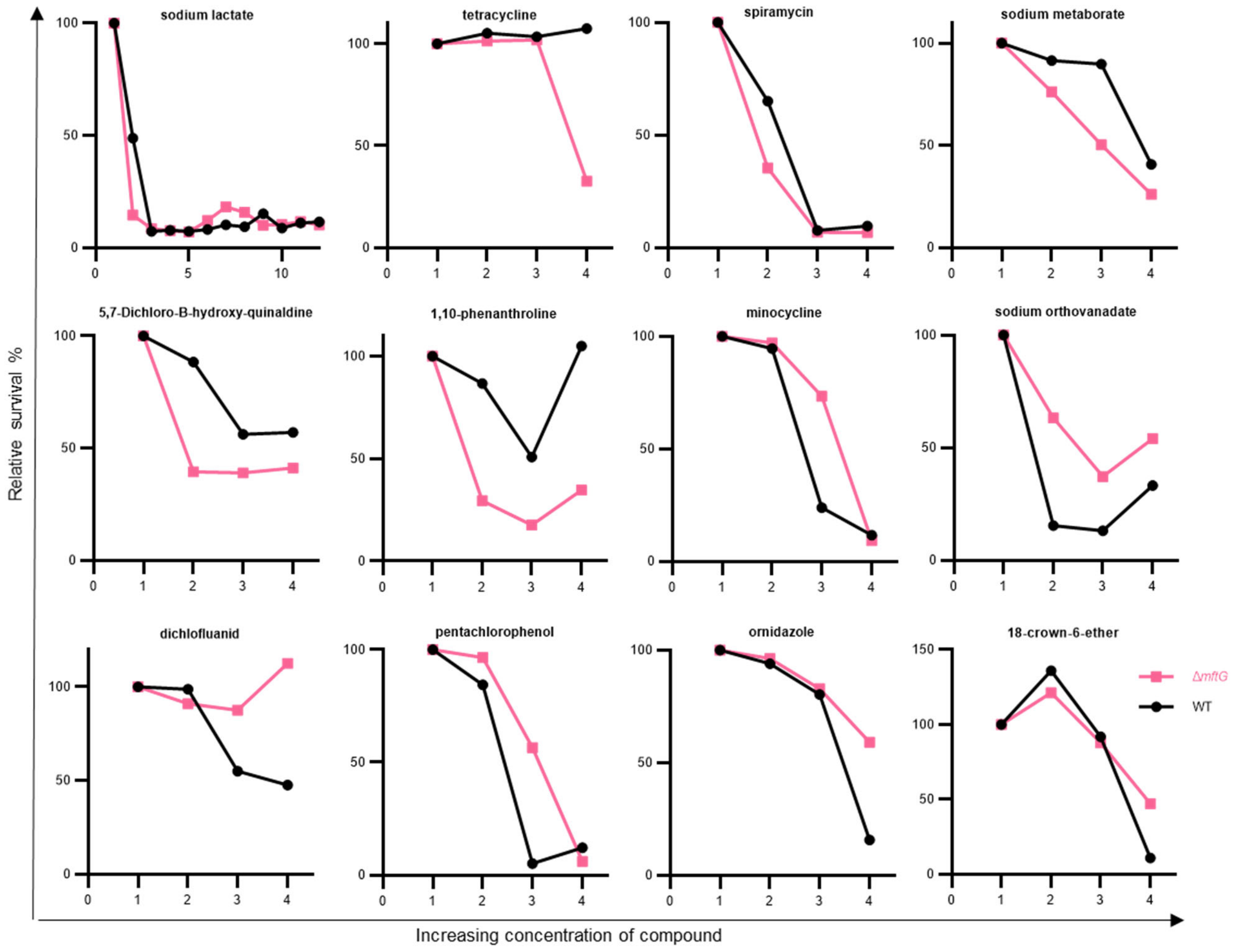


**Supplementary Figure S3 – Differences between the relative growth of the *M. smegmatis* WT (black) and Δ*mftG* (pink) strains using the phenotypic arrays.** Shown are selected compounds from Biolog PM10-20 that modulated the growth behavior. The x-axis represents the concentrations of compounds as multiples of their initial concentration (1x, 2x, 3x, and 4x). The Y-axis shows the relative survival of each strain compared to the starting condition.

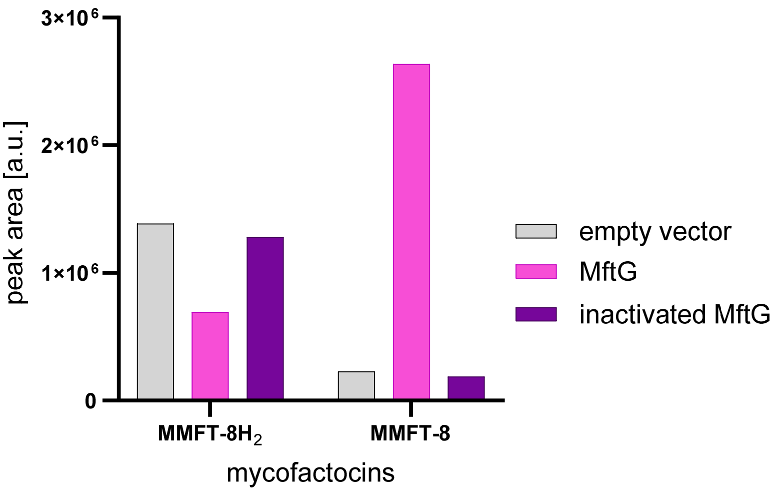


**Supplementary Figure S4 – Mycofactocin dehydrogenase activity of MftG and enzyme preparations obtained from *E. coli* BL21**. Experimental conditions were similar to assays shown in Figure 6E of the manuscript, except that metabolome extract from *M. smegmatis* containing MMFT-8H2 and MMMT-8 was used as a substrate. MftG: enzyme preparation from *E. coli* expressing *mftG*. Empty vector: enzyme preparation obtained from *E. coli* with empty expression vector. Inactivated MftG: enzyme preparation of *E. coli* expressing *mftG*, heat-treated at 99 °C for 10min. Oxidation of MMFT-8$H_2$ to MMFT-8 was associated with the presence of active MftG.

### Supplementary Text 1: Sequences of the plasmids generated in this study.

>pPG17

ATCGGTACCCGGGTTACTTGTACAGCTCGTCCATGCCGTACAGGAACAGGTGGTGGCGGCCCTCGGAGCGCTCGTACTGT
TCCACGATGGTGTAGTCCTCGTTGTGGGAGGTGATGTCCAGCTTGGTGTCCACGTAGTAGTAGCCGGGCAGTTGCACGGG
CTTCTTGGCCATGTAGATGGTCTTGAACTCCACCAGGTAGTGGCCGCCGTCCTTCAGCTTCAGGGCCTGGTGGATCTCGC
CCTTCAGCACGCCGTCGCGGGGGTACAGGCGCTCGGTGGAGGCCTCCCAGCCCATGGTCTTCTTCTGCATTACGGGGCC
GTCGGGGGGGAAGTTGGTGCCGCGCATCTTCACCTTGTAGATCAGCGTGCCGTCCTGCAGGGAGGAGTCCTGGGTCACG
GTCACCAGACCGCCGTCCTCGAAGTTCATCACGCGCTCCCACTTGAAGCCCTCGGGGAAGGACAGCTTCTTGTAATCGGG
GATGTCGGCGGGGTGCTTCACGTACGCCTTGGAGCCGTACATGAACTGGGGGGACAGGATGTCCCAGGCGAAGGGCAGG
GGGCCGCCCTTGGTCACCTTCAGCTTGGCGGTCTGGGTGCCCTCGTAGGGGCGGCCCTCGCCCTCGCCCTCGATCTCGA
ACTCGTGGCCGTTCATGGAGCCCTCCATGCGCACCTTGAAGCGCATGAACTCTTTGATGACCTCCTCGCCCTTGCTCACCA
TTATATATTCCTCCTTTCTGGGATCCTCGGTCGACGTGTGCGCGTAGCCGCTGCGCCAGAACACCCGCATCGCGGTGTCCA
GTGCGACGTCCGGATCGAACTCCTTGTGACGGGCCATGTGCCAATTCTAGATCGTCGAGAAGGTCGACGGCCAGTGGAGG
ACGTACTGCTCCGAGACCTGCTACTGGACCGACGCCGTCGCGTTCCGGGGTGAGTACGAGGGCAGGGAAACCCCGAACA
TGGGACGGCTCACCGGGTTCCGAGAATGGGAGACCCTGCACCACGGCAATCCAATATTGATCCACTAGTCGGTGTCGTAT
GTGCCGAGCGCGGCGATCATCTGCAGGCGGTCGGCGCTCGTGGAAGTGGGCGGGTTCGACGAGACCATGCACTCGGGG
GAGGATGTCGACCTGTGCTGGCGCCTGGTCGAGTCGGGCGCGCGTCTGCGTTACGAACCGATCGCCCTGGTGGCGCACG
ATCACCGCACCAACCTGCGAGCGTGGTTCCACCGCAAGGCATTCTATGGAACGTCGGCCGCTCCGCTGACGGTGCGCCAT
CCCGGTAAGACGTCGCCGCTGGTGATCTCGGGGTGGACGCTGATGGTGTGGCTGATGCTCGGGGTGGGCTCGTTCTTCG
GCTACCTCGCCTCGCTCGCGGCGGCGGTGTTCGCGGGTACCCGCATCGCGCGGGCGCTCAGCGTCGTCGAGACCGAAC
CCAAGGAGGTCGCGGTGGTCGCCGCCCACGGCCTGTGGTCGTCGGCGTTGCAGTTGTGTTCGGCGATCTGTCGCCACTA
CTGGCCCATCGCGATGATCGCGGCGGTGCTGTTCCGCCGGGCGCGGCACGCGGTGCTGGTGGCCGCGGTGGTGGACGG
TGTGGTCGACTGGGTGACGCGACGCGGCAACGCCGACGACGACACCAAACCGGTCGGACTGCTCACCCACATCGTGCTC
AAGCGACTGGACGACATCGCCTACGGCACCGGTCTGTGGACCGGCGTGGTGCGCGAGCGTCACCTCGGCGCGCTCAAGC
CCCAGGTGCGGAGTTAGCCGCCGACTTTGAATTTAAATAACTTCGTATAATGTATGCTATACGAAGTTATCCCAGATCTGTA
AGATAAACTCTAGAAATATTGGATCGTCGGCACCGTCACGGCCGTGGGAGGCGGCACGATCCGCGACGTGATGATCGGCC
GCATCCCCACGGTGCTGCGCAGTGAGCTCTACGCCATCCCGGCGTTGATCTGTGCGTTCGCACGCACAGGCCCGGTGTG
AGAAGGGTCTCTGACGAGCGGGAGAACCCACCCGGGGTGGGCGAGTTTGTCCTGCGTGTGCTCGGTCGAGTAGGCTCTG
GGAGTACCCGTGTGTACGACCAGCACGGCATACATCATTTCGACGCCGAGAGATTCGCCGCCCGAAATGAGCACGATCCG

CATGCTTATAACAGAAAGGAGGTTAATAATGTCGAAGGGCGAGGAGCTGTTCACCGGCGTCGTCCCGATCCTGGTCGAGC
TGGACGGTGACGTCAACGGCCACAAGTTCTCCGTCTCCGGCGAGGGTGAGGGCGACGCCACCTACGGCAAGCTGACCCT
GAAGTTCATCTGCACCACCGGTAAGCTGCCGGTCCCGTGGCCGACCCTGGTCACCACCCTGACCTACGGCGTCCAGTGCT
TCTCCCGCTACCCGGACCACATGAAGCGCCACGACTTCTTCAAGTCCGCCATGCCGGAGGGTTACGTCCAGGAGCGCACC
ATCTCCTTCAAGGACGACGGTAACTACAAGACGCGTGCCGAGGTCAAGTTCGAGGGCGACACCCTGGTCAACCGCATCGA
GCTGAAGGGCATCGACTTCAAGGAGGACGGTAACATCCTGGGCCACAAGCTGGAGTACAACTACAACTCCCACAACGTCT
ACATCACCGCGGACAAGCAGAAGAACGGCATCAAGGCCAACTTCAAGACCCGCCACAACATCGAGGACGGTGGCGTCCA
GCTAGCCGACCACTACCAGCAGAACACCCCGATCGGCGACGGCCCGGTCCTGCTGCCGGACAACCACTACCTGTCCACC
CAGTCCGCCCTGTCCAAGGACCCGAACGAGAAGCGCGACCACATGGTCCTGCTGGAGTTCGTCACCGCCGCCGGCATCA
CCCACGGCATGGACGAGCTGTACAAGTAGATTTATCACCAGCCCGTCATCGTCAACGCCTGATCCGCGGTGCGGACAGGC
CGTGTCGCGACCGGCCGTGCGGAATTAAGCCGGCCCGTACCCTGTGAATAGAGGTCCGCTGTGACACAAGAATCCCTGTT
ACTTCTCGACCGTATTGATTCGGATGATTCCTACGCGAGCCTGCGGAACGACCAGGAATTCTGGGAGCCGCTGGCCCGCC
GAGCCCTGGAGGAGCTCGGGCTGCCGGTGCCGCCGGTGCTGCGGGTGCCCGGCGAGAGCACCAACCCCGTACTGGTCG
GCGAGCCCGACCCGGTGATCAAGCTGTTCGGCGAGCACTGGTGCGGTCCGGAGAGCCTCGCGTCGGAGTCGGAGGCGT
ACGCGGTCCTGGCGGACGCCCCGGTGCCGGTGCCCCGCCTCCTCGGCCGCGGCGAGCTGCGGCCCGGCACCGGAGCC
TGGCCGTGGCCCTACCTGGTGATGAGCCGGATGACCGGCACCACCTGGCGGTCCGCGATGGACGGCACGACCGACCGG
AACGCGCTGCTCGCCCTGGCCCGCGAACTCGGCCGGGTGCTCGGCCGGCTGCACAGGGTGCCGCTGACCGGGAACACC
GTGCTCACCCCCCATTCCGAGGTCTTCCCGGAACTGCTGCGGGAACGCCGCGCGGCGACCGTCGAGGACCACCGCGGGT
GGGGCTACCTCTCGCCCCGGCTGCTGGACCGCCTGGAGGACTGGCTGCCGGACGTGGACACGCTGCTGGCCGGCCGCG
AACCCCGGTTCGTCCACGGCGACCTGCACGGGACCAACATCTTCGTGGACCTGGCCGCGACCGAGGTCACCGGGATCGT
CGACTTCACCGACGTCTATGCGGGAGACTCCCGCTACAGCCTGGTGCAACTGCATCTCAACGCCTTCCGGGGCGACCGCG
AGATCCTGGCCGCGCTGCTCGACGGGGCGCAGTGGAAGCGGACCGAGGACTTCGCCCGCGAACTGCTCGCCTTCACCTT
CCTGCACGACTTCGAGGTGTTCGAGGAGACCCCGCTGGATCTCTCCGGCTTCACCGATCCGGAGGAACTGGCGCAGTTCC
TCTGGGGGCCGCCGGACACCGCCCCCGGCGCCTGACGCCCCGGGCCGCCCGGCGCCGCCCCCGGCCCCCGGCGGCCA
TCGACCCAAGTACCGCCACCTAACAATTCGTTCAAGCCGAGATCGGCTTCCCGGCCGCGGAGTTGTTCGGTAAATTGTCAC
AACGAATTCGATAAAATAAAAAAGGGGACCTCTAGGGTCCCCTTTTTTATTTTATCGAATTCGAGCTCGAAGTTCCTATTCTC
TAGACTGCAGGAACGCGTATAACTTCGTATAATGTATGCTATACGAAGTTATTAATTAATCGACGGCTCGATCATGCCCGCC
ATCACCAGCCGCGGGCCGCACGCGACGATCGCCATGATCGGCCACCGCGCCGCCGAGTTCATCGCGACCTGAGCCGGC
GGATCACCCGCAGCTGTTGCCCGTCGCGCCCGGGTCCCCAGATCCCGGTCAGCACCGCGGCCACCGCGACAACCGAACC
CAGGATGATCACGGCCGAACCGGTCGCGTCCACGAACGCCTGCCGGGCCGTCTGGGCCAGTTCGGCACCCTGCGGTCCC
ATGTGCTCAGCCGCCGTGAGCGCCTCGGCGAGTGAATTGAGCGCGGGTCCGCGTACGGCCTCGGGCATCGCGCCGAGT
GCGGGCGAGAGATGCCTCTCGTACTGCGCTGCGAGCACGGAACCGGCCACCGCGATGCCGAGCGCCGCGCCGATCTCG
CGCGTGGTGTCGTTGACCGCGGACGCGACACCCTGTTTCTCGTCAGGTGCCGCGCCCATGATGGCAGACGTCACCGGTG
CTGTGGACAGCCCGACACCGATGCTCATGATGAACAGTGGCCAGGCGAAATCGAAGTAGGTGGAACCTGTTTCGAGTGTG
CACATGCACCACAACCCCACCGCGGTGACCAGCAGGCCGAGGAATGCCGTGAGGCGCAGGCCGAGCAGCGGCAGGTAC
AGGTGCGTGGTGGCGCCGAACGCCAAGATGGGTACGACCAAGGGGCACAAGGCGATAGCCGTCGCAATCGGGCTGTAAC
CCATGATCAGCTGGATGTACTGCATGCTCACGAAGAAGAACCCGAAGGTCGCGAAGAACAGGAACGTCACCCCCGCCGAG
CCCGTCGCGAAATTCGGGTCGCGGAACAGGCGGACATCGAGCAGCGGGTGGGTGCGCCGCAACTGCACGGCGGTGAAC
GCCGCGGCCAGCGCGACACCGCCCGCCATCGCGCCCCACACCGACGGATGCGTCCACCCGCGCACCGGAGCCTCGACA
ACATGCATGGTTATTGTCTCATGAGCGGATACATATTTGAATGTATTTAGAAAAATGTTTAAACTCGAGGGGGGGCCCGGTG
CTGTATTCGGTTGCAGAGAAACCGGATACACAGCGGACGTTGGGTACCTACGTTGCGTCGAGAGACGAAGGCACCACGAT
TACCGGCGACGATGATCTATTGCCGGTCTACACCGACGCTGGGGTGATTGCGTTTGCTCCGGGAACACCGATCGCTTCGT
TCAAAATCGGCGGAAGATACCCCGTGCGTGCGGTGCTGAAGGACCGCAAAGATGTTCCGTACGGCACCGAGGTCTGCAAG
TACGGGATGAAATCCGGCGAGACCTGTGGGCCGGTCGTTGTGGCTGCCAAGTACACGATCACCGCTCAGCTGCGCACCG
TACATGGCGACTCAGGCGCGCCGCTGTACCGCAAGAACCGCGACGGCACAGCCGATTTGATCGGCATCGCTTCGAGCGTT
AACGAGGACAGTCGCACGACGAAGTTCTTCTGGATCGCGCCCGTGCTGGAAGCACTCAACCTCGAAGCGTGTGGTTGCGG
AGCCATCTAGCAACCACACGAAACATGCGCAACGAACCGCGCAACGAACAACGCCTAGAACTGGCCCTAGATGAGCTGAC
TCGTATCGTTGGTAAACCTAGTTTGACCAGCATGTTTTAACTACGTTCGGTGAGCTGTCAACGGGGCCTGTAACGGCACAA
CGAACCGTGCAACGAGAGTGGCCACGGATGCCACCACAGGCACTACAACGGAGTTCGCCACGTACATCACCACAACCACC
GATTCTGGCGGTGAGCTCCCCGATATTCAGCGGAAATGGCTTGGTATCGACCAAGATTCGTAGAACCCCGTCTCGTCTGGC
TGGTATTCAAAACGGGCGCAACGAAACACGCAACGAGACAGGCATGGCCCAAACCAGAAAACTAGCGTCTACCAGGACTT
TTACCTGTCCGACCCGTTGCAACGGAACCCCCCACGGAACCCCCGCGACACCCGCTCCCCAATTGCGTTAGAACAGCGGT
GGATTGTCGGCTTCGTTGTGGGCCTTTTGAGCCGCTTCCTGTTCTGCCGCACGCTCTTTCCTCGCCCGATAGCCGAGTCGC
TTAACGGTGTCCAGATGCAGCCCGAAATGTTTGGCCGTTTGCGGCCAAGAGTGGCCCTCGTCGTCGTGATAGGCGCGGAT
GCGTTCGCGGCGTGCAGCCTGCTCGGCGAGCCACTCGCTGCGTTCCTGCGCCACGAGCCGGACGACGTGGCGTTCGGAT
AGTCCGGTGATTCGAGCGCCTTCGGCGGCGGTCACGCGCCGCTTTTTGCGGACAGTCGGCTGCCGGTTGTAGCCGTCGC
TGTAGCCGTCGCTGTAGCCGTCGCTCATAGCAATGCCTCCATGGCTGACGCGGACTTTGCGCGCCGCGCAACTGTGCTCG
CCGCCGTGCGCGCTGCTGCGCCCTTCCGCGAGATGGCCGACTGGCGCGCACTGAGTGTGGCCTCGTAGACCACGATCCC
GTCCGCCCAAATGCGCGACTTGGTTGTGATCCAACGCCAAATGCTGTTGGCGATGGCGCGGACCTCGCTGTCCGGTAGCG
GTCCGGGACACACGTCGTTGCACGGGAATTCGGCGTTTCGCGCGTGGCACTCGGCATAGATCGCGCGGCCGAGTCCGTC
CACGTTCCGGGTCGGCAGGTAGATCCGCATGAGGGCGGGACGATAGGCCCACAACCTGACGGAATCGAACAGTGCGCAA
TTCCGCCCTAGCGGCGTCGGAGCCGCTTTGTACGTGGTCTGCTGACGCCAGCGCGGCGGTGGCATGTTCGCGCCGAGCT
CGGCCTCGATGTGGCTGAGTGTGTAGAGATCTGAGTGGAGCCATTCCGTTTCCCAGGCGATGTGGCCGGGGTTTTTGGTC
ATGAGGCCTGAGTAACTGCGGTCGCCGTCGACGGCGCGCCGAAGGCCTTCGGCGCACGCCGCCATGTATGCGAGCGGCT
TACGCCGCGCGTATTCGGTGCGTGGAACAGGGGCGTTGAGTGCCCACACTGCGTGTGCGTGGCCGTTGGCGCGATTGCC
CACGATCGCGTTGGGCAGCGGATGGGACCCCCGGGCGCTGAGCGCTCGGAGCGCTGCGTCTGGATGGTCTACGTCCAC
GACCAGCAGGTTTGCCAGCGCTGTTGGGTTCGCCTCGATGTACCGGCGGCCTAGGGCCGACGCGCGGCTTTGGCGGTAG
ATCCCCTCGAGCAGATCGTCGCTTGCCAGCGGCCAGTACGGCAGCCAGAGCTGCTCAAATTCGTCGGCGACGTGGCTCAC
GCTTGGTAGTAGACCACGATTAATCACCGGTGTATGGTCCGACACGAGCTCCAAGTCAGATATTTCGCTGAGGGGCCACCC
CACAACTGCACACTCCCCCGCTCTCCCGTCGAGCCCTGGTGGTGGAACACCAGCGACAGCCGAGCACCCCCAACCACCT
GTACCAACCAGGAGGAACACATGCGTCGTTTCGAGGACGTTTCCGGGCCGCTGAGAGCCGCTGTGGCGGCCGTACACGC
CGCCTTAGACCCGTTAGACCCCCTGCCGCCTGAATGCGCGGGTACGAGCCACACAGCGCCCGAACTTACGGAGCTGGTG

GGCTCACCTGGCTTTATGGCGTACGAATCGGCTGTGTGCGACCTGTTGGGCGAGGTGAGGTACGCGCTACTCACGCTGGC
AAGGGCGACACAGCCGCCCCACCGAGCCCGCACGGCCGCGCGGTGTCAACAACCGGGTGAGTCGTGCACACCAGCA
GGTGTTCGAGGCTTGGCTCGAAGTGCAGGACATCGTGGCGAACGCCGCCCGATGAGCCGCGCCTTACGCTGGCTGCCAG
CCGTTCGCGGGCTGGTTGGTGCAGCGCGTCGAGCGGTTAGAGGCCCTGCGGTGTTCCACCACCGCAGGCCTCGCCCTTT
TTAAGGCTGAATTTGCTTGTCTCCGAATCCAACTGGCTTGTCCAAGGGTGTATCTACGCTTAATCCAAAGTTCAAACGAGGG
GATTACACATGACCAACTTCGATAACGTTCTCGGCTCGATACTCGTTCCACTGAGCGTCAGACCCCGTAGAAAAGATCAAA
GGATCTTCTTGAGATCCTTTTTTTCTGCGCGTAATCTGCTGCTTGCAAACAAAAAAACCACCGCTACCAGCGGTGGTTTGTT
TGCCGGATCAAGAGCTACCAACTCTTTTTCCGAAGGTAACTGGCTTCAGCAGAGCGCAGATACCAAATACTGTCCTTCTAGT
GTAGCCGTAGTTAGGCCACCACTTCAAGAACTCTGTAGCACCGCCTACATACCTCGCTCTGCTAATCCTGTTACCAGTGGC
TGCTGCCAGTGGCGATAAGTCGTGTCTTACCGGGTTGGACTCAAGACGATAGTTACCGGATAAGGCGCAGCGGTCGGGCT
GAACGGGGGGTTCGTGCACACAGCCCAGCTTGGAGCGAACGACCTACACCGAACTGAGATACCTACAGCGTGAGCTATGA
GAAAGCGCCACGCTTCCCGAAGGGAGAAAGGCGGACAGGTATCCGGTAAGCGGCAGGGTCGGAACAGGAGAGCGCACG
AGGGAGCTTCCAGGGGGAAACGCCTGGTATCTTTATAGTCCTGTCGGGTTTCGCCACCTCTGACTTGAGCGTCGATTTTTG
TGATGCTCGTCAGGGGGGCGGAGCCTATGGAAAAACGCCAGCAACGCGGCCTTTTTACGGTTCCTGGCCTTTTGCTGGCC
TTTTGCTCACATGTTCTTTCCTGCGTTATCCCCTGATTCTGTGGATAACCGTATTACCGCCTTTGAGTGAGCTGATACCGCTC
GCCGCAGCCGAACGACCGAGCGCAGCGAGCGGCCGCTTATTTGTTAACTGTTAATTGTCCTTGTTCAAGGATGCTGTCTTT
GACAACAGATGTTTTCTTGCCTTTGATGTTCAGCAGGAAGCTTGGCGCAAACGTTGATTGTTTGTCTGCGTAGAATCCTCTG
TTTGTCATATAGCTTGTAATCACGACATTGTTTCCTTTCGCTTGAGGTACAGCGAAGTGTGAGTAAGTAAAGGTTACATCGTT
AGGATCAAGATCCATTTTTAACACAAGGCCAGTTTTGTTCAGCGGCTTGTATGGGCCAGTTAAAGAATTAGAAACATAACCA
AGCATGTAAATATCGTTAGACGTAATGCCGTCAATCGTCATTTTTGATCCGCGGGAGTCAGTGAACAGGTACCATTTGCCGT
TCATTTTAAAGACGTTCGCGCGTTCAATTTCATCTGTTACTGTGTTAGATGCAATCAGCGGTTTCATCACTTTTTTCAGTGTG
TAATCATCGTTTAGCTCAATCATACCGAGAGCGCCGTTTGCTAACTCAGCCGTGCGTTTTTTATCGCTTTGCAGAAGTTTTTG
ACTTTCTTGACGGAAGAATGATGTGCTTTTGCCATAGTATGCTTTGTTAAATAAAGATTCTTCGCCTTGGTAGCCATCTTCAG
TTCCAGTGTTTGCTTCAAATACTAAGTATTTGTGGCCTTTATCTTCTACGTAGTGAGGATCTCTCAGCGTATGGTTGTCGCCT
GAGCTGTAGTTGCCTTCATCGATGAACTGCTGTACATTTTGATACGTTTTTCCGTCACCGTCAAAGATTGATTTATAATCCTC
TACACCGTTGATGTTCAAAGAGCTGTCTGATGCTGATACGTTAACTTGTGCAGTTGTCAGTGTTTGTTTGCCGTAATGTTTAC
CGGAGAAATCAGTGTAGAATAAACGGATTTTTCCGTCAGATGTAAATGTGGCTGAACCTGACCATTCTTGTGTTTGGTCTTTT
AGGATAGAATCATTTGCATCGAATTTGTCGCTGTCTTTAAAGACGCGGCCAGCGTTTTTCCAGCTGTCAATAGAAGTTTCGC
CGACTTTTTGATAGAACATGTAAATCGATGTGTCATCCGCATTTTTAGGATCTCCGGCTAATGCAAAGACGATGTGGTAGCC
GTGATAGTTTGCGACAGTGCCGTCAGCGTTTTGTAATGGCCAGCTGTCCCAAACGTCCAGGCCTTTTGCAGAAGAGATATT
TTTAATTGTGGACGAATCGAATTCAGGAACTTGATATTTTTCATTTTTTTGCTGTTCAGGGATTTGCAGCATATCATGGCGTG
TAATATGGGAAATGCCGTATGTTTCCTTATATGGCTTTTGGTTCGTTTCTTTCGCAAACGCTTGAGTTGCGCCTCCTGCCAG
CAGTGCGGTAGTAAAGGTTAATACTGTTGCTTGTTTTGCAAACTTTTTGATGTTCATCGTTCATGTCTCCTTTTTTATGTACTG
TGTTAGCGGTCTGCTTCTTCCAGCCCTCCTGTTTGAAGATGGCAAGTTAGTTACGCACAATAAAAAAAGACCTAAAATATGT
AAGGGGTGACGCCAAAGTATACACTTTGCCCTTTACACATTTTAGGTCTTGCCTGCTTTATCAGTAACAAACCCGCGCGATT
TACTTTTCGACCTCATTCTATTAGACTCTCGTTTGGATTGCAACTGGTCTATTTTCCTCTTTTGTTTGATAGAAAATCATAAAA
GGATTTGCAGACTACGGGCCTAAAGAACTAAAAAATCTATCTGTTTCTTTTCATTCTCTGTATTTTTTATAGTTTCTGTTGCAT
GGGCATAAAGTTGCCTTTTTAATCACAATTCAGAAAATATCATAATATCTCATTTCACTAAATAATAGTGAACGGCAGGTATAT
GGTCGAGCAAGACGTTTCCCGTTGAATATGGCTCATAACACCCCTTGTATTACTGTTTATGTAAGCAGACAGTTTTATTGTTC
ATGATGATATATTTTTATCTTGTGCAATGTAACATCAGAGATTTTGAGACACAACGTGGCTTTCCCCCCCCCCCCCCGACGT
CAGGTGGCTAGAAATGTATCCTAAATCAGAT

>pPG23

GGCCGCGGTACCAGATCTTTAAATCTAGATATCCATGGTCGGACATCTCTCACACCCCCTCTTCCATTCTGGCACTCGATGC
CATATATTTGCGATCTCGATCACAACTGTCGAGACCATACGCGACGAAAGGGAGTCCACATGGAACCGAATCAGCACGTCG
AGGCCGAGACCGAACTCGTCACCGAGACTCTCGTGGAAGAGGTCTCGATCGACGGTATGTGCGGGGTCTACTGACCGTGT
CCACTCAGGTCGCCGAACAGATCACCTTCGACCCGGATGTGAGCTGGCGTCTGCACCATCAGGTCGCGGTTCGGCCGGA
GCCGTTCGGCGCCCTGCTGTACCACTTCGGGACGCGCAAGCTGTCGTTCCTGAAGAACCGAACGGTCGTCGAGGTGGTC
AACTCGCTGGCTGACCACCCCGACGCCCGGTCGGCGCTGTGCGCCGCGGGGGTCGCCGACGATCAGCAAGCGCCCTAT
CTGCACGCACTTGGCGTGCTCGTGCAGTCGAACATGCTCGTACCCGGGAATCCCGAAGGATCGCAATGACTTCAGTTCAG
CCCGTGCCCCGGTTGGTCGAACAGTTCGAGCGCGGACTCGACGCGCCGATCTGCCTCACCTGGGAGCTCACCTACGCGT
GCAACCTCGCGTGTGTCCACTGCCTGTCCTCGTCGGGCAAGCGCGACCCGCGGGAGTTGTCGACGCAGCAGTGCAAGGA
CATCATCGACGAGCTCGAACGCATGCAGGTGTTCTACGTGAACATCGGCGGCGGCGAGCCCACCGTGCGCTCGGACTTCT
GGGAGCTGGTCGACTACGCCACCGCACACCACGTCGGAGTCAAGTTCTCCACCAACGGCGTTCGGATCACGCCCGAGGT
CGCCGCGAAGCTCGCGGCCAGTGACTACGTCGACGTCCAGATCTCGCTGGACGGCGCCAACGCCGAGGTGAACGACGCC
GTGCGCGGCAAGGGCTCGTTCGACATGGCCGTGCGCGCGCTGGAGAACCTGAGCAACGCAGGCTTCACCGACGCCAAGA
TCTCGGTGGTCGTCACGCGGCAGAACGTCGACCAGCTCGACGAATTCGCCGCGCTGGCAGCACGTTACGGCGCCACACT
GCGCATCACGCGGCTGCGGCCGTCCGGTCGGGGCGCCGATGTGTGGGACGATCTGCATCCCACCGCCGAACAGCAGCG
CCAGCTGTACGACTGGCTCGTGGCCAAGGGTGACCGTGTGCTCACCGGCGACTCGTTCTTCCACCTGTCCGGGCTCGGC
GCACCGGGTGCGCTCGCGGGTCTGAACCTGTGCGGCGCAGGCCGCGTGGTGTGCCTGATCGACCCGGTCGGCGACGTC
TACGCCTGCCCGTTCGCGATCCACGACAAGTTCCTGGCCGGAAACATCCTGTCCGACGGTGGTTTCCAGAACGTCTGGCA
GCACTCCGAGCTGTTCCGCGAACTGCGCGAACCGCAGTCGGCCGGTGCCTGCGCAAGCTGCGGGCACTTCGACGCCTGC
CGCGGTGGCTGCATGGCCGCCAAGTTCTTCACGGGGCTGCCGCTCGACGGCCCCGACCCCGAATGCGTCGAGGGCTGG
GGCGCCCCGGCACTGGAGAAGGAACGCGTCAAACCCAAGCCCAGCGGTGACCACTCGCGGGGCACCAAACAAGGGCCG
GTGGCGTTGAAGCTGCTGACCAAACCCCCGGCCCGGTTCTGCAACGAAAGTCCGGTGTGAACATGGCACGAGACATCTGG
TTCGAGACCGTTGCGATCGCCCAGCAAAGGGCCCGCAAGCGGCTTCCGAAATCCGTCTATTCCTCGCTCATCTCCGCGAG
CGAGAAGGGCGTGACGGTCACCGACAACGTCGAGTCGTTCGCCGAGCTCGGTTTCGCACCGCACGTGGTGGGTGCGCCC
GAGAAGCGCGACATGGCCACCACCGTGATGGGCCAACAGATTTCGTTGCCGGTCATCATCTCGCCGACGGGCGTGCAGG
CCGTGCACCCCGACGGCGAGGTCGCGGTTGCTCGGGCCGCCGCCGCGCGCGGTACCGCTATGGGTCTGTCCTCGTTTG
CCAGCAAGCCGATCGAAGAGGTCGTCGCGGTCAACGACAAGATCTTCTTCCAGATCTACTGGCTGGGTGACCGTGACGCG
ATCCTCGCTCGTGCCGAGCGCGCGAAGGCAGCGGGTGCCGTCGGTCTGATCGTCACGACCGACTGGAGCTTCAGTCACG

GCCGTGACTGGGGGAGTCCCAAGATCCCCGAGAAGATGGATCTGAAGACGATGGTCACGATGATGCCCGAGGCGCTGAC
CAAGCCGCGCTGGCTGTGGCAGTGGGGCAAGACCATGCGTCCGCCGAACCTTCGGGTGCCCAACCAGGGGGCGCGCGG
CGAGGACGGCCCGCCGTTCTTTCAGGCCTACGGCGAGTGGATGGGCACGCCCCCGCCGACGTGGGAGGACATCGCCTG
GCTGCGCGAGCAGTGGGACGGGCCGTTCATGCTCAAGGGCGTCATCCGCGTCGACGACGCCAAACGTGCTGTTGACGCG
GGCGTTTCGGCGATCTCGGTGTCCAATCACGGTGGCAACAACCTGGACGGCACACCCGCGGCCATTCGCGCCCTGCCGG
TGATCGCCGAAGCGGTCGGCGACCAGGTCGAGGTATTGCTCGACGGTGGCATCCGCCGCGGCAGCGATGTCGTCAAGGC
TGTCGCGCTGGGTGCGCGTGCGGTGATGATCGGACGCGCCTACCTGTGGGGTCTGGCCGCCGAGGGGCAGGTCGGGGT
CGAGAATGTGCTCGACATCCTGCGCGGAGGTATCGACTCGGCGCTGATGGGCCTGGGGCGTTCGTCGATCCACGATCTG
GTTCCCGAGGACATCCTCGTGCCGGAGGGCTTCACCCGCGCGCTGGGTGTACCGCCCGCATCCGGTTCGTGAGGTCCTC
CCGCGGGGCGTGACAGGGGTGGCAGAAGGGGCGCACGGGATGCGCCCCCTCTTGCGCACGGAAACTGACGGGAAATCG
AACACTGCGTATGCGCGAACATCTGGCGCACGCCAGGTGAATTCGGCTTACCATCGGCACGTGGCTTTTCCCAGCGGGCT
CGGGACCTCGACGTCGAGGCAGCTACACAGCATGGTGCCGATGGTGCTTGTCCCGGTTGGTTCCACCGAACAGCATGGTC
CGCACCTGCCGCTGGACACCGACACCAGGATCGCGGCAGCTGTGGCCGGCACCGTGGTCGAGCAGTTCGGCGCGCCCG
CCGACCGCGACGCGGTGGTGGCCCCTCCGGTGGCCTACGGCGCCAGCGGCGAGCACGAGGGATTTCCCGGAACCGTCT
CGATCGGCACCGCCGCGCTGGAGCTTTTGCTGGTCGAATACGGCAGGTCGGCGTCGAAGTGGACGTCTCGTATCGTCTTC
GTCAACGGTCACGGCGGTAACGTCGAGGCCCTCGCCGCGGCCGTCGCGCTGCTGCGCTACGAGGGCCGCGACGCCGGC
TGGGTGCCGTGCAGCGTGCCCGACGCCGATGCGCACGCGGGCCACACCGAAACGTCCGTACTGCTACATATTTCACCCG
ACGATGTATTGACCGACGAGCTCGTGTGCGGTAATACGGCCCCGTTGGCGGAGCTGATGCCGCGCATGCGCAGTGGCGG
TGTCGCGGCCGTCAGCGAACTCGGGATTCTGGGTGACCCGACGACCGCGACAGCCGCCGAAGGGGAGCGTATCTTTGCT
GAGATGGTCAACGGCTGCGCCGACCGGATCAAGCGGTGGCAGCCCGACCGGAACGGATTGCTGACATGACCGGACCGAG
ACTGCCCGACGGTTTTGCCGTGCAGGTGGATCGCCGAGTCAAGGTGCTCGGAGAGGGCGCGGCGTTGCTGGGCGGCTCA
CCGACCCGCCTGCTGCGGTTGGCGCCGACGGCCCAGAACATGCTCAGCGGGGGCCGCCTCGAGGTTCACGACGCGGTG
TCCGCGCAACTCGCGCGAACCCTCCTGGACGCGACCGTCGCGCATCCGCGGCCTGCCAGCGGGCCGTCCCATCTGGAC
GTCACAGTGGTTGTCCCCGTACGGGACAACGCATCCGGCCTGCACCGCCTGATGGCGGCGCTGCGCGGGCTGCGCGTGA
TCGTGGTCGACGACGGCTCGGCGATCCCGGTGCAACCGTCCGACTTCTCCGGTATGCACTGCGACGTGCAGGTGCTGCG
GCACACCCGCAGCAACGGCCCCGCGGCCGCGCGTAACACCGGCCTTGCGTCGTGCGAGACGGATTTCGTGGCGTTCCTC
GATTCCGACGTGGTGCCCAAGCGTGGTTGGCTCGAGGCGCTGCTCGGGCATTTCTGCGATCCGGCCGTGGCACTGGTGG
CGCCCCGCATCGTGGGTCTGCACAACGCCGACAACATCGTGGCCCGGTACGAGTCCGTCCGATCCTCGCTGGACCTCGG
GGTGCGCGAGGCGCCCGTGGTGCCGCACGGCACGGTGTCGTATGTGCCGAGCGCGGCGATCATCTGCAGGCGGTCGGC
GCTCGTGGAAGTGGGCGGGTTCGACGAGACCATGCACTCGGGGGAGGATGTCGACCTGTGCTGGCGCCTGGTCGAGTCG
GGCGCGCGTCTGCGTTACGAACCGATCGCCCTGGTGGCGCACGATCACCGCACCAACCTGCGAGCGTGGTTCCACCGCA
AGGCATTCTATGGAACGTCGGCCGCTCCGCTGACGGTGCGCCATCCCGGTAAGACGTCGCCGCTGGTGATCTCGGGGTG
GACGCTGATGGTGTGGCTGATGCTCGGGGTGGGCTCGTTCTTCGGCTACCTCGCCTCGCTCGCGGCGGCGGTGTTCGCG
GGTACCCGCATCGCGCGGGCGCTCAGCGTCGTCGAGACCGAACCCAAGGAGGTCGCGGTGGTCGCCGCCCACGGCCTG
TGGTCGTCGGCGTTGCAGTTGTGTTCGGCGATCTGTCGCCACTACTGGCCCATCGCGATGATCGCGGCGGTGCTGTTCCG
CCGGGCGCGGCACGCGGTGCTGGTGGCCGCGGTGGTGGACGGTGTGGTCGACTGGGTGACGCGACGCGGCAACGCCG
ACGACGACACCAAACCGGTCGGACTGCTCACCCACATCGTGCTCAAGCGACTGGACGACATCGCCTACGGCACCGGTCTG
TGGACCGGCGTGGTGCGCGAGCGTCACCTCGGCGCGCTCAAGCCCCAGGTGCGGAGTTAGCCGCCGACTTTGAAAGCTT
ATCGATGTCGACGTAGTTAACTAGCGTACGATCGACTGCCAGGCATCAAATAAAACGAAAGGCTCAGTCGAAAGACTGGGC
CTTTCGTTTTATCTGTTGTTTGTCCGGCCATCATGGCCGCGGTGATCAGCTAGAGCCGTGAACGACAGGGCGAACGCCAG
CCCGCCGACGGCGAGGGTTCCGACCGCTGCAACTCCCGGTGCAACCTTGTCCCGGTCTATTCTCTTCACTGCACCAGCTC
CAATCTGGTGTGAATGCCCCTCGTCTGTTCGCGCAGGCGGGGGGCTCTATTCGTTTGTCAGCATCGAAAGTAGCCAGATCA
GGGATGCGTTGCAACCGCGTATGCCCAGGTCAGAAGAGTCGCACAAGAGTTGCAGACCCCTGGAAAGAAAAATGGCCAGA
GGGCGAAAACACCCTCTGACCAGCGGAGCGGGCGACGGGAATCGAACCCGCGTAGCTAGTTTGGAAGAATGGGTGTCTG
CCGACCACATATATGGGCCGGTCAAGATAGGTTTTTACCCCCTCTCGGCTGCATCCTCTAAGTGGAAAGAAATTGCAGGTC
GTAGAAGCGCGTTGAAGCCTGAGAGTTGCACAGGAGTTGCAACCCGGTAGCCTTGTTCACGACGAGAGGAGACCTAGTTG
GCACGTCGCGGATGGGGATCGCTGAAGACTCAGCGCAGCGGGAGGATCCAAGCCTCATACGTCAACCCGCAGGACGGTG
TGAGGTACTACGCGCTGCAGACCTACGACAACAAGATGGACGCCGAAGCCTGGCTCGCGGGCGAGAAGCGGCTCATCGA
GATGGAGACCTGGACCCCTCCACAGGACCGGGCGAAGAAGGCAGCCGCCAGCGCCATCACGCTGGAGGAGTACACCCG
GAAGTGGCTCGTGGAGCGCGACCTCGCAGACGGCACCAGGGATCTGTACAGCGGGCACGCGGAGCGCCGCATCTACCC
GGTGCTAGGTGAAGTGGCGGTCACAGAGATGACGCCAGCTCTGGTGCGTGCGTGGTGGGCCGGGATGGGTAGGAAGCA
CCCGACTGCCCGCCGGCATGCCTACAACGTCCTCCGGGCGGTGATGAACACAGCGGTCGAGGACAAGCTGATCGCAGAG
AACCCGTGCCGGATCGAGCAGAAGGCAGCCGATGAGCGCGACGTAGAGGCGCTGACGCCTGAGGAGCTGGACATCGTC
GCCGCTGAGATCTTCGAGCACTACCGGATCGCGGCATACATCCTGGCGTGGACGAGCCTCCGGTTCGGAGAGCTGATCG
AGCTTCGCCGCAAGGACATCGTGGACGACGGCATGACGATGAAGCTCCGGGTGCGCCGTGGCGCTTCCCGCGTGGGGAA
CAAGATCGTCGTTGGCAACGCCAAGACCGTCCGGTCGAAGCGTCCTGTGACGGTTCCGCCTCACGTCGCGGAGATGATCC
GAGCGCACATGAAGGACCGTACGAAGATGAACAAGGGCCCCGAGGCATTCCTGGTGACCACGACGCAGGGCAACCGGCT
GTCGAAGTCCGCGTTCACCAAGTCGCTGAAGCGTGGCTACGCCAAGATCGGTCGGCCGGAACTCCGCATCCACGACCTCC
GCGCTGTCGGCGCTACGTTCGCCGCTCAGGCAGGTGCGACGACCAAGGAGCTGATGGCCCGTCTCGGTCACACGACTCC
TAGGATGGCGATGAAGTACCAGATGGCGTCTGAGGCCCGCGACGAGGCTATCGCTGAGGCGATGTCCAAGCTGGCCAAG
ACCTCCTGAAACGCAAAAAGCCCCCCTCCCAAGGACACTGAGTCCTAAAGAGGGGGGTTTCTTGTCAGTACGCGAAGAAC
CACGCCTGGCCGCGAGCGCCAGCACCGCCGCTCTGTGCGGAGACCTGGGCACCAGCCCCGCCGCCGCCAGGAGCATTG
CCGTTCCCGCCAGCTAGCAACAAAGCGACGTTGTGTCTCAAAATCTCTGATGTTACATTGCACAAGATAAAAATATATCATC
ATGAACAATAAAACTGTCTGCTTACATAAACAGTAATACAAGGGGTGTTATGAGCCATATTCAACGGGAAACGTCTTGCTCG
AGGCCGCGATTAAATTCCAACATGGATGCTGATTTATATGGGTATAAATGGGCTCGCGATAATGTCGGGCAATCAGGTGCG
ACAATCTATCGCTTGTATGGGAAGCCCCATGCGCCAGAGTTGTTTCTGAAACATGGCAAAGGTAGCGTTGCCAATGATGTT
ACAGATGAGATGGTCAGACTAAACTGGCTGACGGAATTTATGCCTCTTCCGACCATCAAGCATTTTATCCGTACTCCTGATG
ATGCATGGTTACTCACCACTGCGATCCCCGGGAAAACAGCATTCCAGGTATTAGAAGAATATCCTGATTCAGGTGAAAATAT
TGTTGATGCGCTGGCAGTGTTCCTGCGCCGGTTGCATTCGATTCCTGTTTGTAATTGTCCTTTTAACAGCGATCGCGTATTT
CGTCTCGCTCAGGCGCAATCACGAATGAATAACGGTTTGGTTGATGCGAGTGATTTTGATGACGAGCGTAATGGCTGGCCT
GTTGAACAAGTCTGGAAAGAAATGCATAATCTTTTGCCATTCTCACCGGATTCAGTCGTCACTCATGGTGATTTCTCACTTGA
TAACCTTATTTTTGACGAGGGGAAATTAATAGGTTGTATTGATGTTGGACGAGTCGGAATCGCAGACCGATACCAGGATCTT

GCCATCCTATGGAACTGCCTCGGTGAGTTTTCTCCTTCATTACAGAAACGGCTTTTTCAAAAATATGGTATTGATAATCCTGA
TATGAATAAATTGCAGTTTCATTTGATGCTCGATGAGTTTTTCTAATCAGAATTGGTTAATTGGTTGTAACACTGGCAGAGCA
TTACGCTGACTTGACGGGACGGCGGCTTTGTTGAATAAATCGAACTTTTGCTGAGTTGAAGGATCAGATCACGCATCTTCCC
GACAACGCAGACCGTTCCGTGGCAAAGCAAAAGTTCAAAATCACCAACTGGTCCACCTACAACAAAGCTCTCATCAACCGT
GGCTCCCTCACTTTCTGGCTGGATGATGGGGCGATTCAGGCCTGGTATGAGTCAGCAACACCTTCTTCACGAGGCAGACC
TCACTAGTTCCACTGAGCGTCAGACCCCGTAGAAAAGATCAAAGGATCTTCTTGAGATCCTTTTTTTCTGCGCGTAATCTGC
TGCTTGCAAACAAAAAAACCACCGCTACCAGCGGTGGTTTGTTTGCCGGATCAAGAGCTACCAACTCTTTTTCCGAAGGTAA
CTGGCTTCAGCAGAGCGCAGATACCAAATACTGTCCTTCTAGTGTAGCCGTAGTTAGGCCACCACTTCAAGAACTCTGTAG
CACCGCCTACATACCTCGCTCTGCTAATCCTGTTACCAGTGGCTGCTGCCAGTGGCGATAAGTCGTGTCTTACCGGGTTGG
ACTCAAGACGATAGTTACCGGATAAGGCGCAGCGGTCGGGCTGAACGGGGGGTTCGTGCACACAGCCCAGCTTGGAGCG
AACGACCTACACCGAACTGAGATACCTACAGCGTGAGCATTGAGAAAGCGCCACGCTTCCCGAAGGGAGAAAGGCGGACA
GGTATCCGGTAAGCGGCAGGGTCGGAACAGGAGAGCGCACGAGGGAGCTTCCAGGGGGAAACGCCTGGTATCTTTATAG
TCCTGTCGGGTTTCGCCACCTCTGACTTGAGCGTCGATTTTTGTGATGCTCGTCAGGGGGGCGGAGCCTATGGAAAAACG
CCAGCAACGCGGCCTTTTTACGGTTCCTGGCCTTTTGCTGGCCTTTTGCTCACATGTTCTTTCCTGCGTTATCCCCTGATTC
TGTGGATAACCGTATTACCGCCTTTGAGTGAGCTGATACCGCTCGCCGCAGCCGAACGACCGAGCGCAACGCGTGC

>pPG29

GGCCGCGGTACCAGATCTTTAAATCTAGATATCCATGGCTCTCACACCCCCTCTTCCATTCTGGCACTCGATGCCATATATT
TGCGATCTCGATCACAACTGTCGAGACCATACGCATATGGAGTTAGCCGCCGACTTTGAGTGATGTGTTGATCGTCGGCGC
GGGCAGCGCCGGATCGGTGTTGGCCGAACGCCTTTCCGCGGACCCGAGCTGTCAGGTGACGGTGGTGGAGGCCGGCCC
GGCCCCGTCGGATCCGCGGGTGGCCGCGCAGATCACCGACGGCGTGCGGCTACCCATCGGCGCCGCGAGTTCGGTGGT
GCGGCACTATGGGTCGACGCTGACCGAGGATCCGCCGCGGCGCACCGAGATCATCCGCGGTTCGGTGGTCGGCGGCTC
CGGAGCGGTCAACGGTGGCTATTTCTGTCGCGGCCTGCCCACCGACTTCACCGCGTGGGCGGTTCCCGGATGGAGCTGG
GACGATGTGCTGCCCCATTTCCGTGCGATCGAGACCGACCTCGACTTCGCCACCGCGCTGCACGGCTCCGAAGGTCCGAT
CAAGGTGCGCCGGGTCCACGAATTCGACGGCTGCACCGCGGCATTCGTCGACGCCGTATCCGCCGCGGGGTATGGCTGG
ATCGACGATCTCAACGGGTCGGATGCCGACGCGGCGTTGCCACCCGGCGTTGGCGCGGTGCCGCTGAACATCGACGGCG
GAACCCGTCTCGGACCCGGAGGCGCCTACCTGCAGCCCGCACTCGACCGGCCCAATCTGCATCTGCGCGCCGACACCCG
GGTGCGTCGCGTGCTGGTCGAGCATGATGCCGCGGTGGGTGTGGAATGCGTCGACGGTGAGATCCTGTACGCAGATCGA
ATCGTGTTGTCCGCCGGGGCCATCGGCTCGGCACATCTGTTGCTGCTCTCAGGCATCGGGCCGGCGGGCGACTTGGCGG
CCCACGGGATCGCGGTGGCGGCGAACCTACCTGTGGGGACGGCGACAGTCGACCACCCGGAGTGGGTGCTGCCGGTGG
CCTGGACCCCCACACATGACCTGCCGCCGCTGGAAGCCGTGCTCACCACGGCCGACGGCATCGAGATCCGGCCCTACAC
CGCGGGTTTCTCGGCGCTGGTGCACGGGCCCGAACACGACCCGGCCGAACTCCCACATCTCGGTGTCGCGCTCATGCGT
CCGCATTCACGGGGCAGGGTGCGTCTCGCGTCGGCGGATCCGGCCGTGCCCCCGATCATCGAACACCGTTACGACACGG
TCGCAGGGGATGTCGACGCGCTGCGCGCGGGCGCCGAACTCGCCCGCGAATTGGTAAGTCACGCAGTCAAAGTGGGGG
AGGCCTCCTGGTCGACTTCGCAGCACCTCGCGGGGACCGCGCCGCTGAGTGCCGACGGGCAAGGCGTGCTGGACCCGA
CGTGCCGGGTACTCGGTGTCGAGCGGTTGTGGGTGGTCGACGGCTCGATCATGCCCGCCATCACCAGCCGCGGGCCGCA
CGCGACGATCGCCATGATCGGCCACCGCGCCGCCGAGTTCATCGCGACCTGAGCCGGCGGATCACCCGCAGCTGTTAAG
CTTATCGATGTCGACGTAGTTAACTAGCGTACGATCGACTGCCAGGCATCAAATAAAACGAAAGGCTCAGTCGAAAGACTG
GGCCTTTCGTTTTATCTGTTGTTTGTCCGGCCATCATGGCCGCGGTGATCAGCTAGAGCCGTGAACGACAGGGCGAACGC
CAGCCCGCCGACGGCGAGGGTTCCGACCGCTGCAACTCCCGGTGCAACCTTGTCCCGGTCTATTCTCTTCACTGCACCAG
CTCCAATCTGGTGTGAATGCCCCTCGTCTGTTCGCGCAGGCGGGGGGCTCTATTCGTTTGTCAGCATCGAAAGTAGCCAG
ATCAGGGATGCGTTGCAACCGCGTATGCCCAGGTCAGAAGAGTCGCACAAGAGTTGCAGACCCCTGGAAAGAAAAATGGC
CAGAGGGCGAAAACACCCTCTGACCAGCGGAGCGGGCGACGGGAATCGAACCCGCGTAGCTAGTTTGGAAGAATGGGTG
TCTGCCGACCACATATATGGGCCGGTCAAGATAGGTTTTTACCCCCTCTCGGCTGCATCCTCTAAGTGGAAAGAAATTGCA
GGTCGTAGAAGCGCGTTGAAGCCTGAGAGTTGCACAGGAGTTGCAACCCGGTAGCCTTGTTCACGACGAGAGGAGACCTA
GTTGGCACGTCGCGGATGGGGATCGCTGAAGACTCAGCGCAGCGGGAGGATCCAAGCCTCATACGTCAACCCGCAGGAC
GGTGTGAGGTACTACGCGCTGCAGACCTACGACAACAAGATGGACGCCGAAGCCTGGCTCGCGGGCGAGAAGCGGCTCA
TCGAGATGGAGACCTGGACCCCTCCACAGGACCGGGCGAAGAAGGCAGCCGCCAGCGCCATCACGCTGGAGGAGTACAC
CCGGAAGTGGCTCGTGGAGCGCGACCTCGCAGACGGCACCAGGGATCTGTACAGCGGGCACGCGGAGCGCCGCATCTA
CCCGGTGCTAGGTGAAGTGGCGGTCACAGAGATGACGCCAGCTCTGGTGCGTGCGTGGTGGGCCGGGATGGGTAGGAA
GCACCCGACTGCCCGCCGGCATGCCTACAACGTCCTCCGGGCGGTGATGAACACAGCGGTCGAGGACAAGCTGATCGCA
GAGAACCCGTGCCGGATCGAGCAGAAGGCAGCCGATGAGCGCGACGTAGAGGCGCTGACGCCTGAGGAGCTGGACATC
GTCGCCGCTGAGATCTTCGAGCACTACCGGATCGCGGCATACATCCTGGCGTGGACGAGCCTCCGGTTCGGAGAGCTGAT
CGAGCTTCGCCGCAAGGACATCGTGGACGACGGCATGACGATGAAGCTCCGGGTGCGCCGTGGCGCTTCCCGCGTGGG
GAACAAGATCGTCGTTGGCAACGCCAAGACCGTCCGGTCGAAGCGTCCTGTGACGGTTCCGCCTCACGTCGCGGAGATGA
TCCGAGCGCACATGAAGGACCGTACGAAGATGAACAAGGGCCCCGAGGCATTCCTGGTGACCACGACGCAGGGCAACCG
GCTGTCGAAGTCCGCGTTCACCAAGTCGCTGAAGCGTGGCTACGCCAAGATCGGTCGGCCGGAACTCCGCATCCACGAC
CTCCGCGCTGTCGGCGCTACGTTCGCCGCTCAGGCAGGTGCGACGACCAAGGAGCTGATGGCCCGTCTCGGTCACACGA
CTCCTAGGATGGCGATGAAGTACCAGATGGCGTCTGAGGCCCGCGACGAGGCTATCGCTGAGGCGATGTCCAAGCTGGC
CAAGACCTCCTGAAACGCAAAAAGCCCCCCTCCCAAGGACACTGAGTCCTAAAGAGGGGGGTTTCTTGTCAGTACGCGAA
GAACCACGCCTGGCCGCGAGCGCCAGCACCGCCGCTCTGTGCGGAGACCTGGGCACCAGCCCCGCCGCCGCCAGGAGC
ATTGCCGTTCCCGCCAGCTAGCAACAAAGCGACGTTGTGTCTCAAAATCTCTGATGTTACATTGCACAAGATAAAAATATAT
CATCATGAACAATAAAACTGTCTGCTTACATAAACAGTAATACAAGGGGTGTTATGAGCCATATTCAACGGGAAACGTCTTG
CTCGAGGCCGCGATTAAATTCCAACATGGATGCTGATTTATATGGGTATAAATGGGCTCGCGATAATGTCGGGCAATCAGG
TGCGACAATCTATCGCTTGTATGGGAAGCCCCATGCGCCAGAGTTGTTTCTGAAACATGGCAAAGGTAGCGTTGCCAATGA
TGTTACAGATGAGATGGTCAGACTAAACTGGCTGACGGAATTTATGCCTCTTCCGACCATCAAGCATTTTATCCGTACTCCT
GATGATGCATGGTTACTCACCACTGCGATCCCCGGGAAAACAGCATTCCAGGTATTAGAAGAATATCCTGATTCAGGTGAA
AATATTGTTGATGCGCTGGCAGTGTTCCTGCGCCGGTTGCATTCGATTCCTGTTTGTAATTGTCCTTTTAACAGCGATCGCG
TATTTCGTCTCGCTCAGGCGCAATCACGAATGAATAACGGTTTGGTTGATGCGAGTGATTTTGATGACGAGCGTAATGGCT
GGCCTGTTGAACAAGTCTGGAAAGAAATGCATAATCTTTTGCCATTCTCACCGGATTCAGTCGTCACTCATGGTGATTTCTC
ACTTGATAACCTTATTTTTGACGAGGGGAAATTAATAGGTTGTATTGATGTTGGACGAGTCGGAATCGCAGACCGATACCAG

GATCTTGCCATCCTATGGAACTGCCTCGGTGAGTTTTCTCCTTCATTACAGAAACGGCTTTTTCAAAAATATGGTATTGATAA
TCCTGATATGAATAAATTGCAGTTTCATTTGATGCTCGATGAGTTTTTCTAATCAGAATTGGTTAATTGGTTGTAACACTGGCA
GAGCATTACGCTGACTTGACGGGACGGCGGCTTTGTTGAATAAATCGAACTTTTGCTGAGTTGAAGGATCAGATCACGCAT
CTTCCCGACAACGCAGACCGTTCCGTGGCAAAGCAAAAGTTCAAAATCACCAACTGGTCCACCTACAACAAAGCTCTCATC
AACCGTGGCTCCCTCACTTTCTGGCTGGATGATGGGGCGATTCAGGCCTGGTATGAGTCAGCAACACCTTCTTCACGAGG
CAGACCTCACTAGTTCCACTGAGCGTCAGACCCCGTAGAAAAGATCAAAGGATCTTCTTGAGATCCTTTTTTTCTGCGCGTA
ATCTGCTGCTTGCAAACAAAAAAACCACCGCTACCAGCGGTGGTTTGTTTGCCGGATCAAGAGCTACCAACTCTTTTTCCGA
AGGTAACTGGCTTCAGCAGAGCGCAGATACCAAATACTGTCCTTCTAGTGTAGCCGTAGTTAGGCCACCACTTCAAGAACT
CTGTAGCACCGCCTACATACCTCGCTCTGCTAATCCTGTTACCAGTGGCTGCTGCCAGTGGCGATAAGTCGTGTCTTACCG
GGTTGGACTCAAGACGATAGTTACCGGATAAGGCGCAGCGGTCGGGCTGAACGGGGGGTTCGTGCACACAGCCCAGCTT
GGAGCGAACGACCTACACCGAACTGAGATACCTACAGCGTGAGCATTGAGAAAGCGCCACGCTTCCCGAAGGGAGAAAGG
CGGACAGGTATCCGGTAAGCGGCAGGGTCGGAACAGGAGAGCGCACGAGGGAGCTTCCAGGGGGAAACGCCTGGTATCT
TTATAGTCCTGTCGGGTTTCGCCACCTCTGACTTGAGCGTCGATTTTTGTGATGCTCGTCAGGGGGGCGGAGCCTATGGAA
AAACGCCAGCAACGCGGCCTTTTTACGGTTCCTGGCCTTTTGCTGGCCTTTTGCTCACATGTTCTTTCCTGCGTTATCCCCT
GATTCTGTGGATAACCGTATTACCGCCTTTGAGTGAGCTGATACCGCTCGCCGCAGCCGAACGACCGAGCGCAACGCGTG
C

>pPG32

TGTCTCAAAATCTCTGATGTTACATTGCACAAGATAAAAATATATCATCATGAACAATAAAACTGTCTGCTTACATAAACAGTA
ATACAAGGGGTGTTATGAGCCATATTCAACGGGAAACGTCTTGCTCGAGGCCGCGATTAAATTCCAACATGGATGCTGATTT
ATATGGGTATAAATGGGCTCGCGATAATGTCGGGCAATCAGGTGCGACAATCTATCGCTTGTATGGGAAGCCCCATGCGCC
AGAGTTGTTTCTGAAACATGGCAAAGGTAGCGTTGCCAATGATGTTACAGATGAGATGGTCAGACTAAACTGGCTGACGGA
ATTTATGCCTCTTCCGACCATCAAGCATTTTATCCGTACTCCTGATGATGCATGGTTACTCACCACTGCGATCCCCGGGAAA
ACAGCATTCCAGGTATTAGAAGAATATCCTGATTCAGGTGAAAATATTGTTGATGCGCTGGCAGTGTTCCTGCGCCGGTTGC
ATTCGATTCCTGTTTGTAATTGTCCTTTTAACAGCGATCGCGTATTTCGTCTCGCTCAGGCGCAATCACGAATGAATAACGG
TTTGGTTGATGCGAGTGATTTTGATGACGAGCGTAATGGCTGGCCTGTTGAACAAGTCTGGAAAGAAATGCATAATCTTTTG
CCATTCTCACCGGATTCAGTCGTCACTCATGGTGATTTCTCACTTGATAACCTTATTTTTGACGAGGGGAAATTAATAGGTTG
TATTGATGTTGGACGAGTCGGAATCGCAGACCGATACCAGGATCTTGCCATCCTATGGAACTGCCTCGGTGAGTTTTCTCC
TTCATTACAGAAACGGCTTTTTCAAAAATATGGTATTGATAATCCTGATATGAATAAATTGCAGTTTCATTTGATGCTCGATGA
GTTTTTCTAATCAGAATTGGTTAATTGGTTGTAACACTGGCAGAGCATTACGCTGACTTGACGGGACGGCGGCTTTGTTGAA
TAAATCGAACTTTTGCTGAGTTGAAGGATCAGATCACGCATCTTCCCGACAACGCAGACCGTTCCGTGGCAAAGCAAAAGT
TCAAAATCACCAACTGGTCCACCTACAACAAAGCTCTCATCAACCGTGGCTCCCTCACTTTCTGGCTGGATGATGGGGCGA
TTCAGGCCTGGTATGAGTCAGCAACACCTTCTTCACGAGGCAGACCTCACTAGTTCCACTGAGCGTCAGACCCCGTAGAAA
AGATCAAAGGATCTTCTTGAGATCCTTTTTTTCTGCGCGTAATCTGCTGCTTGCAAACAAAAAAACCACCGCTACCAGCGGT
GGTTTGTTTGCCGGATCAAGAGCTACCAACTCTTTTTCCGAAGGTAACTGGCTTCAGCAGAGCGCAGATACCAAATACTGTC
CTTCTAGTGTAGCCGTAGTTAGGCCACCACTTCAAGAACTCTGTAGCACCGCCTACATACCTCGCTCTGCTAATCCTGTTAC
CAGTGGCTGCTGCCAGTGGCGATAAGTCGTGTCTTACCGGGTTGGACTCAAGACGATAGTTACCGGATAAGGCGCAGCGG
TCGGGCTGAACGGGGGGTTCGTGCACACAGCCCAGCTTGGAGCGAACGACCTACACCGAACTGAGATACCTACAGCGTG
AGCATTGAGAAAGCGCCACGCTTCCCGAAGGGAGAAAGGCGGACAGGTATCCGGTAAGCGGCAGGGTCGGAACAGGAGA
GCGCACGAGGGAGCTTCCAGGGGGAAACGCCTGGTATCTTTATAGTCCTGTCGGGTTTCGCCACCTCTGACTTGAGCGTC
GATTTTTGTGATGCTCGTCAGGGGGGCGGAGCCTATGGAAAAACGCCAGCAACGCGGCCTTTTTACGGTTCCTGGCCTTTT
GCTGGCCTTTTGCTCACATGTTCTTTCCTGCGTTATCCCCTGATTCTGTGGATAACCGTATTACCGCCTTTGAGTGAGCTGA
TACCGCTCGCCGCAGCCGAACGACCGAGCGCAACGCGTGCGGCCGCGGTACCAGATCTTTAAATCTAGATATCCATGGCT
CTCACACCCCCTCTTCCATTCTGGCACTCGATGCCATATATTTGCGATCTCGATCACAACTGTCGAGACCATACGCATATGG
AGTTAGCCGCCGACTTTGAGTGATGTGTTGATCGTCGGCGCGGGCAGCGCCGGATCGGTGTTGGCCGAACGCCTTTCCG
CGGACCCGAGCTGTCAGGTGACGGTGGTGGAGGCCGGCCCGGCCCCGTCGGATCCGCGGGTGGCCGCGCAGATCACCG
ACGGCGTGCGGCTACCCATCGGCGCCGCGAGTTCGGTGGTGCGGCACTATGGGTCGACGCTGACCGAGGATCCGCCGC
GGCGCACCGAGATCATCCGCGGTTCGGTGGTCGGCGGCTCCGGAGCGGTCAACGGTGGCTATTTCTGTCGCGGCCTGCC
CACCGACTTCACCGCGTGGGCGGTTCCCGGATGGAGCTGGGACGATGTGCTGCCCCATTTCCGTGCGATCGAGACCGAC
CTCGACTTCGCCACCGCGCTGCACGGCTCCGAAGGTCCGATCAAGGTGCGCCGGGTCCACGAATTCGACGGCTGCACCG
CGGCATTCGTCGACGCCGTATCCGCCGCGGGGTATGGCTGGATCGACGATCTCAACGGGTCGGATGCCGACGCGGCGTT
GCCACCCGGCGTTGGCGCGGTGCCGCTGAACATCGACGGCGGAACCCGTCTCGGACCCGGAGGCGCCTACCTGCAGCC
CGCACTCGACCGGCCCAATCTGCATCTGCGCGCCGACACCCGGGTGCGTCGCGTGCTGGTCGAGCATGATGCCGCGGTG
GGTGTGGAATGCGTCGACGGTGAGATCCTGTACGCAGATCGAATCGTGTTGTCCGCCGGGGCCATCGGCTCGGCACATCT
GTTGCTGCTCTCAGGCATCGGGCCGGCGGGCGACTTGGCGGCCCACGGGATCGCGGTGGCGGCGAACCTACCTGTGGG
GACGGCGACAGTCGACCACCCGGAGTGGGTGCTGCCGGTGGCCTGGACCCCCACACATGACCTGCCGCCGCTGGAAGC
CGTGCTCACCACGGCCGACGGCATCGAGATCCGGCCCTACACCGCGGGTTTCTCGGCGCTGGTGCACGGGCCCGAACAC
GACCCGGCCGAACTCCCACATCTCGGTGTCGCGCTCATGCGTCCGCATTCACGGGGCAGGGTGCGTCTCGCGTCGGCGG
ATCCGGCCGTGCCCCCGATCATCGAACACCGTTACGACACGGTCGCAGGGGATGTCGACGCGCTGCGCGCGGGCGCCG
AACTCGCCCGCGAATTGGTAAGTCACGCAGTCAAAGTGGGGGAGGCCTCCTGGTCGACTTCGCAGCACCTCGCGGGGAC
CGCGCCGCTGAGTGCCGACGGGCAAGGCGTGCTGGACCCGACGTGCCGGGTACTCGGTGTCGAGCGGTTGTGGGTGGT
CGACGGCTCGATCATGCCCGCCATCACCAGCCGCGGGCCGCACGCGACGATCGCCATGATCGGCCACCGCGCCGCCGA
GTTCATCGCGACCCACCACCACCACCACCACTAATAGTAAGCTTATCGATGTCGACGTAGTTAACTAGCGTACGATCGACT
GCCAGGCATCAAATAAAACGAAAGGCTCAGTCGAAAGACTGGGCCTTTCGTTTTATCTGTTGTTTGTCCGGCCATCATGGC
CGCGGTGATCAGCTAGAGCCGTGAACGACAGGGCGAACGCCAGCCCGCCGACGGCGAGGGTTCCGACCGCTGCAACTC
CCGGTGCAACCTTGTCCCGGTCTATTCTCTTCACTGCACCAGCTCCAATCTGGTGTGAATGCCCCTCGTCTGTTCGCGCAG
GCGGGGGGCTCTATTCGTTTGTCAGCATCGAAAGTAGCCAGATCAGGGATGCGTTGCAACCGCGTATGCCCAGGTCAGAA
GAGTCGCACAAGAGTTGCAGACCCCTGGAAAGAAAAATGGCCAGAGGGCGAAAACACCCTCTGACCAGCGGAGCGGGCG
ACGGGAATCGAACCCGCGTAGCTAGTTTGGAAGAATGGGTGTCTGCCGACCACATATATGGGCCGGTCAAGATAGGTTTTT
ACCCCCTCTCGGCTGCATCCTCTAAGTGGAAAGAAATTGCAGGTCGTAGAAGCGCGTTGAAGCCTGAGAGTTGCACAGGA
GTTGCAACCCGGTAGCCTTGTTCACGACGAGAGGAGACCTAGTTGGCACGTCGCGGATGGGGATCGCTGAAGACTCAGC

GCAGCGGGAGGATCCAAGCCTCATACGTCAACCCGCAGGACGGTGTGAGGTACTACGCGCTGCAGACCTACGACAACAA
GATGGACGCCGAAGCCTGGCTCGCGGGCGAGAAGCGGCTCATCGAGATGGAGACCTGGACCCCTCCACAGGACCGGGC
GAAGAAGGCAGCCGCCAGCGCCATCACGCTGGAGGAGTACACCCGGAAGTGGCTCGTGGAGCGCGACCTCGCAGACGG
CACCAGGGATCTGTACAGCGGGCACGCGGAGCGCCGCATCTACCCGGTGCTAGGTGAAGTGGCGGTCACAGAGATGACG
CCAGCTCTGGTGCGTGCGTGGTGGGCCGGGATGGGTAGGAAGCACCCGACTGCCCGCCGGCATGCCTACAACGTCCTCC
GGGCGGTGATGAACACAGCGGTCGAGGACAAGCTGATCGCAGAGAACCCGTGCCGGATCGAGCAGAAGGCAGCCGATGA
GCGCGACGTAGAGGCGCTGACGCCTGAGGAGCTGGACATCGTCGCCGCTGAGATCTTCGAGCACTACCGGATCGCGGCA
TACATCCTGGCGTGGACGAGCCTCCGGTTCGGAGAGCTGATCGAGCTTCGCCGCAAGGACATCGTGGACGACGGCATGA
CGATGAAGCTCCGGGTGCGCCGTGGCGCTTCCCGCGTGGGGAACAAGATCGTCGTTGGCAACGCCAAGACCGTCCGGTC
GAAGCGTCCTGTGACGGTTCCGCCTCACGTCGCGGAGATGATCCGAGCGCACATGAAGGACCGTACGAAGATGAACAAG
GGCCCCGAGGCATTCCTGGTGACCACGACGCAGGGCAACCGGCTGTCGAAGTCCGCGTTCACCAAGTCGCTGAAGCGTG
GCTACGCCAAGATCGGTCGGCCGGAACTCCGCATCCACGACCTCCGCGCTGTCGGCGCTACGTTCGCCGCTCAGGCAGG
TGCGACGACCAAGGAGCTGATGGCCCGTCTCGGTCACACGACTCCTAGGATGGCGATGAAGTACCAGATGGCGTCTGAG
GCCCGCGACGAGGCTATCGCTGAGGCGATGTCCAAGCTGGCCAAGACCTCCTGAAACGCAAAAAGCCCCCCTCCCAAGG
ACACTGAGTCCTAAAGAGGGGGGTTTCTTGTCAGTACGCGAAGAACCACGCCTGGCCGCGAGCGCCAGCACCGCCGCTC
TGTGCGGAGACCTGGGCACCAGCCCCGCCGCCGCCAGGAGCATTGCCGTTCCCGCCAGCTAGCAACAAAGCGACGTTG

>pPG36_biocat

CCGACACCATCGAATGGTGCAAAACCTTTCGCGGTATGGCATGATAGCGCCCGGAAGAGAGTCAATTCAGGGTGGTGAAT
GTGAAACCAGTAACGTTATACGATGTCGCAGAGTATGCCGGTGTCTCTTATCAGACCGTTTCCCGCGTGGTGAACCAGGCC
AGCCACGTTTCTGCGAAAACGCGGGAAAAAGTGGAAGCGGCGATGGCGGAGCTGAATTACATTCCCAACCGCGTGGCACA
ACAACTGGCGGGCAAACAGTCGTTGCTGATTGGCGTTGCCACCTCCAGTCTGGCCCTGCACGCGCCGTCGCAAATTGTCG
CGGCGATTAAATCTCGCGCCGATCAACTGGGTGCCAGCGTGGTGGTGTCGATGGTAGAACGAAGCGGCGTCGAAGCCTG
TAAAGCGGCGGTGCACAATCTTCTCGCGCAACGCGTCAGTGGGCTGATCATTAACTATCCGCTGGATGACCAGGATGCCAT
TGCTGTGGAAGCTGCCTGCACTAATGTTCCGGCGTTATTTCTTGATGTCTCTGACCAGACACCCATCAACAGTATTATTTTCT
CCCATGAAGACGGTACGCGACTGGGCGTGGAGCATCTGGTCGCATTGGGTCACCAGCAAATCGCGCTGTTAGCGGGCCC
ATTAAGTTCTGTCTCGGCGCGTCTGCGTCTGGCTGGCTGGCATAAATATCTCACTCGCAATCAAATTCAGCCGATAGCGGA
ACGGGAAGGCGACTGGAGTGCCATGTCCGGTTTTCAACAAACCATGCAAATGCTGAATGAGGGCATCGTTCCCACTGCGA
TGCTGGTTGCCAACGATCAGATGGCGCTGGGCGCAATGCGCGCCATTACCGAGTCCGGGCTGCGCGTTGGTGCGGATAT
CTCGGTAGTGGGATACGACGATACCGAAGACAGCTCATGTTATATCCCGCCGTTAACCACCATCAAACAGGATTTTCGCCT
GCTGGGGCAAACCAGCGTGGACCGCTTGCTGCAACTCTCTCAGGGCCAGGCGGTGAAGGGCAATCAGCTGTTGCCCGTC
TCACTGGTGAAAAGAAAAACCACCCTGGCGCCCAATACGCAAACCGCCTCTCCCCGCGCGTTGGCCGATTCATTAATGCAG
CTGGCACGACAGGTTTCCCGACTGGAAAGCGGGCAGTGAGCGCAACGCAATTAATGTAAGTTAGCTCACTCATTAGGCACA
ATTCTCATGTTTGACAGCTTATCATCGACTGCACGGTGCACCAATGCTTCTGGCGTCAGGCAGCCATCGGAAGCTGTGGTA
TGGCTGTGCAGGTCGTAAATCACTGCATAATTCGTGTCGCTCAAGGCGCACTCCCGTTCTGGATAATGTTTTTTGCGCCGA
CATCATAACGGTTCTGGCAAATATTCTGAAATGAGCTGTTGACAATTAATCATCGGCTCGTATAATGTGTGGAATTGTGAGC
GGATAACAATTTCACACAGGAAACAGCCAGTCCGTTTAGGTGTTTTCACGAGCACTTCACCAACAAGGACCATAGCATATGA
AAATCGAAGAAGGTAAACTGGTAATCTGGATTAACGGCGATAAAGGCTATAACGGTCTCGCTGAAGTCGGTAAGAAATTCG
AGAAAGATACCGGAATTAAAGTCACCGTTGAGCATCCGGATAAACTGGAAGAGAAATTCCCACAGGTTGCGGCAACTGGCG
ATGGCCCTGACATTATCTTCTGGGCACACGACCGCTTTGGTGGCTACGCTCAATCTGGCCTGTTGGCTGAAATCACCCCGG
ACAAAGCGTTCCAGGACAAGCTGTATCCGTTTACCTGGGATGCCGTACGTTACAACGGCAAGCTGATTGCTTACCCGATCG
CTGTTGAAGCGTTATCGCTGATTTATAACAAAGATCTGCTGCCGAACCCGCCAAAAACCTGGGAAGAGATCCCGGCGCTGG
ATAAAGAACTGAAAGCGAAAGGTAAGAGCGCGCTGATGTTCAACCTGCAAGAACCGTACTTCACCTGGCCGCTGATTGCTG
CTGACGGGGGTTATGCGTTCAAGTATGAAAACGGCAAGTACGACATTAAAGACGTGGGCGTGGATAACGCTGGCGCGAAA
GCGGGTCTGACCTTCCTGGTTGACCTGATTAAAAACAAACACATGAATGCAGACACCGATTACTCCATCGCAGAAGCTGCC
TTTAATAAAGGCGAAACAGCGATGACCATCAACGGCCCGTGGGCATGGTCCAACATCGACACCAGCAAAGTGAATTATGGT
GTAACGGTACTGCCGACCTTCAAGGGTCAACCATCCAAACCGTTCGTTGGCGTGCTGAGCGCAGGTATTAACGCCGCCAG
TCCGAACAAAGAGCTGGCAAAAGAGTTCCTCGAAAACTATCTGCTGACTGATGAAGGTCTGGAAGCGGTTAATAAAGACAA
ACCGCTGGGTGCCGTAGCGCTGAAGTCTTACGAGGAAGAGTTGGTGAAAGATCCGCGGATTGCCGCCACTATGGAAAACG
CCCAGAAAGGTGAAATCATGCCGAACATCCCGCAGATGTCCGCTTTCTGGTATGCCGTGCGTACTGCGGTGATCAACGCC
GCCAGCGGTCGTCAGACTGTCGATGAAGCCCTGAAAGACGCGCAGACTAATTCGAGCTCGAACAACAACAACAATAACAAT
AACAACAACCTCGGGATCGAGGGAAGGATTTCAGAATTCGTTCTGATTGTTGGTGCCGGCAGCGCCGGTAGCGTGCTGGC
AGAACGTCTGAGCGCCGATCCGAGCTGTCAGGTGACCGTGGTTGAAGCAGGTCCGGCACCGAGTGATCCGCGCGTGGCA
GCACAGATTACCGATGGCGTGCGCCTGCCGATTGGTGCCGCCAGTAGCGTGGTTCGTCATTATGGCAGCACCCTGACCGA
AGATCCGCCGCGTCGCACCGAAATTATTCGCGGTAGTGTGGTTGGTGGCAGTGGTGCAGTTAATGGCGGTTATTTTTGTCG
TGGTCTGCCGACCGATTTTACCGCCTGGGCAGTTCCGGGCTGGAGCTGGGATGATGTTCTGCCGCATTTTCGTGCCATTG
AAACCGATCTGGATTTTGCCACCGCCCTGCATGGCAGCGAAGGTCCGATTAAGGTTCGTCGCGTGCATGAATTTGATGGTT
GTACCGCAGCATTTGTTGATGCAGTTAGTGCCGCAGGTTATGGCTGGATTGATGATCTGAATGGCAGTGATGCAGATGCAG
CACTGCCGCCGGGCGTGGGTGCTGTTCCTCTGAATATTGATGGCGGCACCCGTCTGGGTCCGGGTGGTGCATATCTGCAG
CCGGCACTGGATCGTCCGAATCTGCATCTGCGCGCAGATACCCGTGTTCGTCGCGTTCTGGTGGAACATGATGCAGCCGT
TGGTGTGGAATGCGTGGATGGTGAAATTCTGTATGCAGATCGTATTGTGCTGAGTGCCGGCGCAATTGGCAGCGCCCATCT
GCTGCTGCTGAGCGGCATTGGTCCGGCAGGTGACCTGGCCGCACATGGCATTGCCGTGGCAGCCAATCTGCCGGTTGGC
ACCGCCACCGTTGATCATCCGGAATGGGTTCTGCCGGTGGCATGGACCCCGACCCATGATCTGCCGCCGCTGGAAGCCG
TTCTGACCACCGCCGATGGCATTGAAATTCGTCCGTATACCGCCGGTTTTAGCGCCCTGGTGCATGGTCCGGAACATGATC
CGGCAGAACTGCCGCATCTGGGTGTGGCCCTGATGCGTCCGCATAGTCGCGGCCGCGTTCGCCTGGCAAGTGCAGATCC
GGCAGTTCCGCCGATTATTGAACATCGCTATGATACCGTTGCAGGCGATGTTGATGCCCTGCGTGCCGGTGCAGAACTGG
CCCGTGAACTGGTGAGTCATGCAGTGAAAGTGGGTGAAGCCAGCTGGAGCACCAGTCAGCATCTGGCCGGTACCGCCCC
GCTGAGTGCAGATGGCCAGGGTGTGCTGGATCCGACCTGCCGCGTTCTGGGTGTGGAACGTCTGTGGGTTGTGGATGGC
AGCATTATGCCGGCCATTACCAGCCGTGGCCCGCATGCAACCATTGCAATGATTGGCCATCGTGCCGCAGAGTTTATTGCC
ACCTAAAAGCTTGGCACTGGCCGTCGTTTTACAACGTCGTGACTGGGAAAACCCTGGCGTTACCCAACTTAATCGCCTTGC
AGCACATCCCCCTTTCGCCAGCTGGCGTAATAGCGAAGAGGCCCGCACCGATCGCCCTTCCCAACAGTTGCGCAGCCTGA

ATGGCGAATGGCAGCTTGGCTGTTTTGGCGGATGAGATAAGATTTTCAGCCTGATACAGATTAAATCAGAACGCAGAAGCG
GTCTGATAAAACAGAATTTGCCTGGCGGCAGTAGCGCGGTGGTCCCACCTGACCCCATGCCGAACTCAGAAGTGAAACGC
CGTAGCGCCGATGGTAGTGTGGGGTCTCCCCATGCGAGAGTAGGGAACTGCCAGGCATCAAATAAAACGAAAGGCTCAGT
CGAAAGACTGGGCCTTTCGTTTTATCTGTTGTTTGTCGGTGAACGCTCTCCTGAGTAGGACAAATCCGCCGGGAGCGGATT
TGAACGTTGCGAAGCAACGGCCCGGAGGGTGGCGGGCAGGACGCCCGCCATAAACTGCCAGGCATCAAATTAAGCAGAA
GGCCATCCTGACGGATGGCCTTTTTGCGTTTCTACAAACTCTTTTGTTTATTTTTCTAAATACATTCAAATATGTATCCGCTCA
TGAGACAATAACCCTGATAAATGCTTCAATAATATTGAAAAAGGAAGAGTATGAGTATTCAACATTTCCGTGTCGCCCTTATT
CCCTTTTTTGCGGCATTTTGCCTTCCTGTTTTTGCTCACCCAGAAACGCTGGTGAAAGTAAAAGATGCTGAAGATCAGTTGG
GTGCACGAGTGGGTTACATCGAACTGGATCTCAACAGCGGTAAGATCCTTGAGAGTTTTCGCCCCGAAGAACGTTTCCCAA
TGATGAGCACTTTTAAAGTTCTGCTATGTGGCGCGGTATTATCCCGTGTTGACGCCGGGCAAGAGCAACTCGGTCGCCGCA
TACACTATTCTCAGAATGACTTGGTTGAGTACTCACCAGTCACAGAAAAGCATCTTACGGATGGCATGACAGTAAGAGAATT
ATGCAGTGCTGCCATAACCATGAGTGATAACACTGCGGCCAACTTACTTCTGACAACGATCGGAGGACCGAAGGAGCTAAC
CGCTTTTTTGCACAACATGGGGGATCATGTAACTCGCCTTGATCGTTGGGAACCGGAGCTGAATGAAGCCATACCAAACGA
CGAGCGTGACACCACGATGCCTGTAGCAATGGCAACAACGTTGCGCAAACTATTAACTGGCGAACTACTTACTCTAGCTTC
CCGGCAACAATTAATAGACTGGATGGAGGCGGATAAAGTTGCAGGACCACTTCTGCGCTCGGCCCTTCCGGCTGGCTGGT
TTATTGCTGATAAATCTGGAGCCGGTGAGCGTGGGTCTCGCGGTATCATTGCAGCACTGGGGCCAGATGGTAAGCCCTCC
CGTATCGTAGTTATCTACACGACGGGGAGTCAGGCAACTATGGATGAACGAAATAGACAGATCGCTGAGATAGGTGCCTCA
CTGATTAAGCATTGGTAACTGTCAGACCAAGTTTACTCATATATACTTTAGATTGATTTACCCCGGTTGATAATCAGAAAAGC
CCCAAAAACAGGAAGATTGTATAAGCAAATATTTAAATTGTAAACGTTAATATTTTGTTAAAATTCGCGTTAAATTTTTGTTAAA
TCAGCTCATTTTTTAACCAATAGGCCGAAATCGGCAAAATCCCTTATAAATCAAAAGAATAGACCGAGATAGGGTTGAGTGT
TGTTCCAGTTTGGAACAAGAGTCCACTATTAAAGAACGTGGACTCCAACGTCAAAGGGCGAAAAACCGTCTATCAGGGCGA
TGGCCCACTACGTGAACCATCACCCAAATCAAGTTTTTTGGGGTCGAGGTGCCGTAAAGCACTAAATCGGAACCCTAAAGG
GAGCCCCCGATTTAGAGCTTGACGGGGAAAGCCGGCGAACGTGGCGAGAAAGGAAGGGAAGAAAGCGAAAGGAGCGGG
CGCTAGGGCGCTGGCAAGTGTAGCGGTCACGCTGCGCGTAACCACCACACCCGCCGCGCTTAATGCGCCGCTACAGGGC
GCGTAAAAGGATCTAGGTGAAGATCCTTTTTGATAATCTCATGACCAAAATCCCTTAACGTGAGTTTTCGTTCCACTGAGCG
TCAGACCCCGTAGAAAAGATCAAAGGATCTTCTTGAGATCCTTTTTTTCTGCGCGTAATCTGCTGCTTGCAAACAAAAAAAC
CACCGCTACCAGCGGTGGTTTGTTTGCCGGATCAAGAGCTACCAACTCTTTTTCCGAAGGTAACTGGCTTCAGCAGAGCGC
AGATACCAAATACTGTCCTTCTAGTGTAGCCGTAGTTAGGCCACCACTTCAAGAACTCTGTAGCACCGCCTACATACCTCGC
TCTGCTAATCCTGTTACCAGTGGCTGCTGCCAGTGGCGATAAGTCGTGTCTTACCGGGTTGGACTCAAGACGATAGTTACC
GGATAAGGCGCAGCGGTCGGGCTGAACGGGGGGTTCGTGCACACAGCCCAGCTTGGAGCGAACGACCTACACCGAACTG
AGATACCTACAGCGTGAGCTATGAGAAAGCGCCACGCTTCCCGAAGGGAGAAAGGCGGACAGGTATCCGGTAAGCGGCA
GGGTCGGAACAGGAGAGCGCACGAGGGAGCTTCCAGGGGGAAACGCCTGGTATCTTTATAGTCCTGTCGGGTTTCGCCA
CCTCTGACTTGAGCGTCGATTTTTGTGATGCTCGTCAGGGGGGCGGAGCCTATGGAAAAACGCCAGCAACGCGGCCTTTT
TACGGTTCCTGGCCTTTTGCTGGCCTTTTGCTCACATGTTCTTTCCTGCGTTATCCCCTGATTCTGTGGATAACCGTATTACC
GCCTTTGAGTGAGCTGATACCGCTCGCCGCAGCCGAACGACCGAGCGCAGCGAGTCAGTGAGCGAGGAAGCGGAAGAGC
GCCTGATGCGGTATTTTCTCCTTACGCATCTGTGCGGTATTTCACACCGCATATATGGTGCACTCTCAGTACAATCTGCTCT
GATGCCGCATAGTTAAGCCAGTATACACTCCGCTATCGCTACGTGACTGGGTCATGGCTGCGCCCCGACACCCGCCAACA
CCCGCTGACGCGCCCTGACGGGCTTGTCTGCTCCCGGCATCCGCTTACAGACAAGCTGTGACCGTCTCCGGGAGCTGCAT
GTGTCAGAGGTTTTCACCGTCATCACCGAAACGCGCGAGGCAGCTGCGGTAAAGCTCATCAGCGTGGTCGTGCAGCGATT
CACAGATGTCTGCCTGTTCATCCGCGTCCAGCTCGTTGAGTTTCTCCAGAAGCGTTAATGTCTGGCTTCTGATAAAGCGGG
CCATGTTAAGGGCGGTTTTTTCCTGTTTGGTCACTGATGCCTCCGTGTAAGGGGGATTTCTGTTCATGGGGGTAATGATACC
GATGAAACGAGAGAGGATGCTCACGATACGGGTTACTGATGATGAACATGCCCGGTTACTGGAACGTTGTGAGGGTAAACA
ACTGGCGGTATGGATGCGGCGGGACCAGAGAAAAATCACTCAGGGTCAATGCCAGCGCTTCGTTAATACAGATGTAGGTG
TTCCACAGGGTAGCCAGCAGCATCCTGCGATGCAGATCCGGAACATAATGGTGCAGGGCGCTGACTTCCGCGTTTCCAGA
CTTTACGAAACACGGAAACCGAAGACCATTCATGTTGTTGCTCAGGTCGCAGACGTTTTGCAGCAGCAGTCGCTTCACGTT
CGCTCGCGTATCGGTGATTCATTCTGCTAACCAGTAAGGCAACCCCGCCAGCCTAGCCGGGTCCTCAACGACAGGAGCAC
GATCATGCGCACCCGTGGCCAGGACCCAACGCTGCCCGAAATT
